# Automatic mechanistic inference from large families of Boolean models generated by Monte Carlo Tree Search

**DOI:** 10.1101/2022.10.13.512151

**Authors:** Bryan Glazer, Jonathan Lifferth, Carlos F. Lopez

## Abstract

1

**Motivation:** Many important processes in biology, such as signaling and gene regulation, can be described using logic models. These logic models are typically built to behaviorally emulate experimentally observed phenotypes, which are assumed to be steady states of a biological system. Most models are built by hand and therefore researchers are only able to consider one or perhaps a few potential mechanisms. We present a method to automatically synthesize Boolean logic models with a specified set of steady states. Our method, called MC-Boomer, is based on Monte Carlo Tree Search (MCTS), an efficient, parallel search method using reinforcement learning. Our approach enables users to constrain the model search space using prior knowledge or biochemical interaction databases, thus leading to generation of biologically plausible mechanistic hypotheses. Our approach can generate very large numbers of data-consistent models. To help develop mechanistic insight from these models, we developed analytical tools for multi-model inference and model selection. These tools reveal the key sets of interactions that govern the behavior of the models.

**Results:** We demonstrate that MC-Boomer works well at reconstructing randomly generated models. Then, using single time point measurements and reasonable biological constraints, our method generates hundreds of thousands of candidate models that match experimentally validated *in-vivo* behaviors of the *Drosophila* segment polarity network. Finally we outline how our multimodel analysis procedures elucidate potentially novel biological mechanisms and provide opportunities for model-driven experimental validation.

**Availability:** Code is available at: www.github.com/bglazer/mcboomer

## 2 Introduction

Technological advances in high throughput sequencing have significantly increased the amount of data available to biologists. However, the systems of molecular interactions that generate many cellular phenotypes remain poorly understood. This lack of understanding is a particularly pressing problem for diseases such as cancer, in which small genetic perturbations can have drastic clinical consequences. In order to understand and potentially intervene in the mechanisms by which cellular systems become dysregulated, one must first create a hypothesis of the system’s interactions.

### 2.1 Computational modeling of biology

Given the complexity and non-linearity of many biological systems, computational models are a key tool for hypothesis generation and testing, allowing *in-silico* perturbation and experimentation. Much previous work has shown the value of computational models of cellular systems for both understanding mechanisms and predicting cellular response to perturbation (Béal *et al*., 2019; Sáez-Rodríguez *et al*., 2007; Schlatter *et al*., 2009).

In this work, we focus on automatic synthesis of Boolean models. Boolean models are a simple, two-state, dynamical system with Boolean logic update rules introduced by Kauffman (1969). Boolean models do not have reaction rate parameters; their behavior is entirely determined by the structure of the update rules. Despite their simplicity, previous work has shown their accuracy and utility in modeling a wide variety of biological systems.

### 2.2 Manually specifying models is difficult

However, manually creating these computational models can be time-consuming and difficult for several reasons. First, selecting a set of interactions that lead to the desired behavior is challenging due to the vast number of possible interactions. Further, introducing a new interaction can create feedback loops that change the model’s behavior in unintuitive ways. Finally, data is often limited, only covering a limited set of conditions. Thus, many possible model configurations may have behavior that matches the (limited) data equally well. In order to have a reasonable chance of finding a model that captures an accurate representation of the biological system, including in conditions outside the given data, one must create many models.

### 2.3 Our approach

Thus, automated model synthesis is desirable as it alleviates the difficulty of manually constructing a wide variety of models that are consistent with data. However, an efficient search algorithm is required to synthesize data-consistent models from the vast space of possible Boolean models. Inspired by recent work in reinforcement learning for games, which also have combinatorially large search spaces, we investigate Monte Carlo Tree Search (MCTS). Our method uses MCTS to iteratively build Boolean models by adding molecular interactions to the model’s update rules, similar to the way this algorithm is used to select moves in the games of chess or Go (Gelly *et al*., 2006).

We show that MCTS works well for a wide variety of input data and model structures by testing the algorithm’s ability to recover randomly generated Boolean models. Further, we show that it works for a more biologically realistic scenario: generating multi-cellular models of the *Drosophila* segment polarity network. Our method generated hundreds of thousands of models of the segment polarity network that are all consistent with experimental observations.

Having created a large collection of data-consistent models, one must derive some insight into the key interactions or mechanisms which drive their behavior. This is itself a challenging pattern recognition problem, which we address by developing data driven methods to extract mechanisms from models. Specifically, we present methods for clustering models based on the structure of their interactions. Using the structural clustering, our methods reveal the key interactions that control model behavior. We employ this analysis to develop a novel hypothesis for the mechanism of regulation of the *wg* gene by isoforms of *CI* in *Drosophila*.

We call this pipeline of automated model generation and mechanism exploration MC-Boomer, or Monte Carlo Boolean Modeler.

### 2.4 Previous Work

Approaches to inferring logical models with data-consistent behavior can be divided into two categories: constraint solving and optimization. Constraint solving based methods pose the problem as a series of logical constraints, e.g. that the update functions must be consistent with steady states described in the data. These constraints are typically encoded as Boolean logic equations or in a more abstract formalism such as answer set programming (ASP) (Chevalier *et al*., 2020, 2019) or satisfiability modulo theories (SMT) problems (Yordanov *et al*., 2016; Fisher *et al*., 2015). Specialized solvers then find a set of models which satisfy all the constraints specified by the data and the modeling assumptions.

Optimization methods use general purpose discrete optimization algorithms to generate Boolean models, which are then scored according to a user-defined objective function (incorporating e.g. similarity to data or model complexity). The optimization algorithms then generate new models which are variations of the best scoring models (Lim *et al*., 2016; Terfve *et al*., 2012).

### 2.5 Comparison to our method

Our method differs from previous approaches in several key ways. First, we employ a heuristic optimization method, in contrast to linear programming or satisfiability solver based approaches. This allows us to trivially encode more complex model dynamics (e.g. multi-cellularity) and constraints on the form of update rules. Further, our optimization approach requires simulation of all models, giving us a view into the state spaces of our models. This allows us to characterize models according their behavior between initial conditions and steady states, yielding greater insight into populations of models that all have similar steady states. This comes at the cost of greater required computational resources compared to methods based on specialized constraint solvers. However, our method is trivially parallelizable, which exploit to find large numbers of data-consistent models in a reasonable time frame. Finally, our optimization based approach immediately generates models that are partial matches to the experimental data. In contrast, constraint solvers may neglect useful models that do not perfectly satisfy constraints, even when those constraints are misspecified or based on noisy data. In the worst case, constraint solvers may yield zero models after a lengthy search, while our approximation approach yields a spectrum of models of varying complexity and goodness of fit to the data.

MC-Boomer is conceptually similar to optimization based approaches such as BTR (Lim *et al*., 2016) and PRUNET (Rodriguez *et al*., 2015). One novel aspect of our approach is that we employ Monte Carlo Tree Search (MCTS) (Kocsis and Szepesvári, 2006). The efficiency of MCTS allows us to find large numbers of models that fit the data well. Thus, we are able to make inferences about possible mechanisms of biological systems that are based on families of thousands of potential models. Some previous approaches (Saez-Rodriguez *et al*., 2009) consider the relative probabilities of individual interactions, based on the whole population of data-consistent models. However, we investigate model structures with more sophisticated and fine-grained analyses, such as structure-based clustering and cluster interpretability methods.

## 3 Methods

Here we describe the components of our framework for automated generation and exploration of mechanistic hypotheses, which we call MC-Boomer (Monte Carlo Boolean Modeler). As shown in Fig 1, our framework consists of three steps: gathering data and prior knowledge (Fig 1, left), using Monte Carlo Tree Search to generate and test model hypotheses (Fig 1, middle), and finally analyzing the model collection using statistical and multi-model inference approaches (Fig 1, right). In this section, we primarily describe the second step, the algorithmic components involved in generating models. We describe the third step, analysis of the models generated by MCTS, in more detail in the Results (Section 4), as part of our analysis of the segment polarity network.

**Figure 1:**
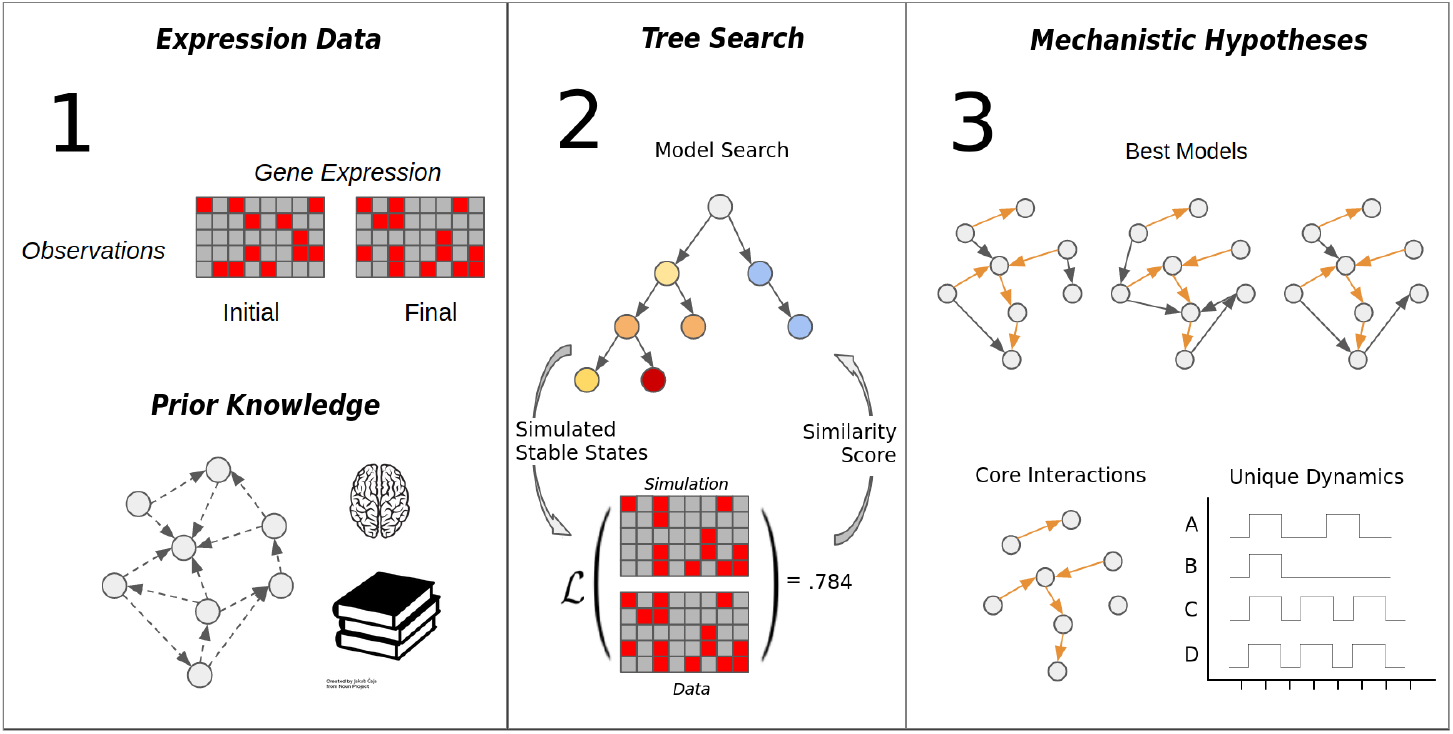
MC-Boomer workflow. Broadly speaking, MC-Boomer consists of three steps. The first is to gather data of the steady state expression levels of the genes of interest in a system. Optionally, one can gather information regarding known interactions between genes in the system. Potential sources for this information include biochemical interaction databases, published literature, or even prior experience and intuition. This prior knowledge can guide and constrain the second step, model generation. We use Monte Carlo Tree Search to generate models (Section 3.3 and Fig. 3). The objective of search is to find models that have simulated attractor states that are similar to data, as measured by an edit distance metric, described in Section 3.2. We test this algorithm’s ability to recover random models in Section 3.4. With promising results on random models, we apply the method to a more biologically realistic model: the *Drosophila* segment polarity network, described in Section 4.1. This resulted in *>* 200*k* high quality models, which we analyze further in Sections 4.1.2 to extract structural features.

We separate our discussion of model generation (Fig 1, middle) into three sections: simulation, scoring, and search. We simulate our models with Boolean update rules, introduce a novel edit distance metric for scoring, and employ Monte Carlo Tree Search (MCTS) for search. Below we will describe each component in more detail.

### 3.1 Simulation

A Boolean model is composed of logic rules that determine the state of each species in the system at the next step. We use synchronous updating, meaning that at each step we apply the corresponding update equations to *every* species in the model. Synchronous updating is deterministic and is guaranteed to reach either a single stable attractor state or a sequence of periodically repeating states, called a cyclic attractor. We detect both stable and cyclic attractors by tracking previous states and halting simulation when the state matches a previously simulated state.

We restrict the form of our Boolean update functions to only dominant inhibition, having the form:

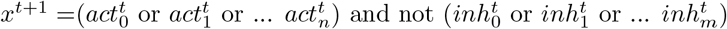

Here, *x* is the species in the model that will be updated, 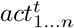 and 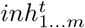 are the states (at time *t*) of other species in the network that regulate the target node. Both *act*_*i*_ and *inh*_*i*_ can be a single species or composites of of two or more species connected by an *and* clause, e.g. (*a* and *b*). A node is activated at *t* + 1 only if one or more of its activators is active and no inhibitors are active at *t*.

A more comprehensive review of simulating biological systems with Boolean networks can be found in Albert *et al*. (2008).

### 3.2 Scoring

We developed an edit distance metric that compares reference data to model steady states (described in Figure 2). This distance is used to guide the MCTS search algorithm towards models that generate steady states that are similar to the data.

**Figure 2:**
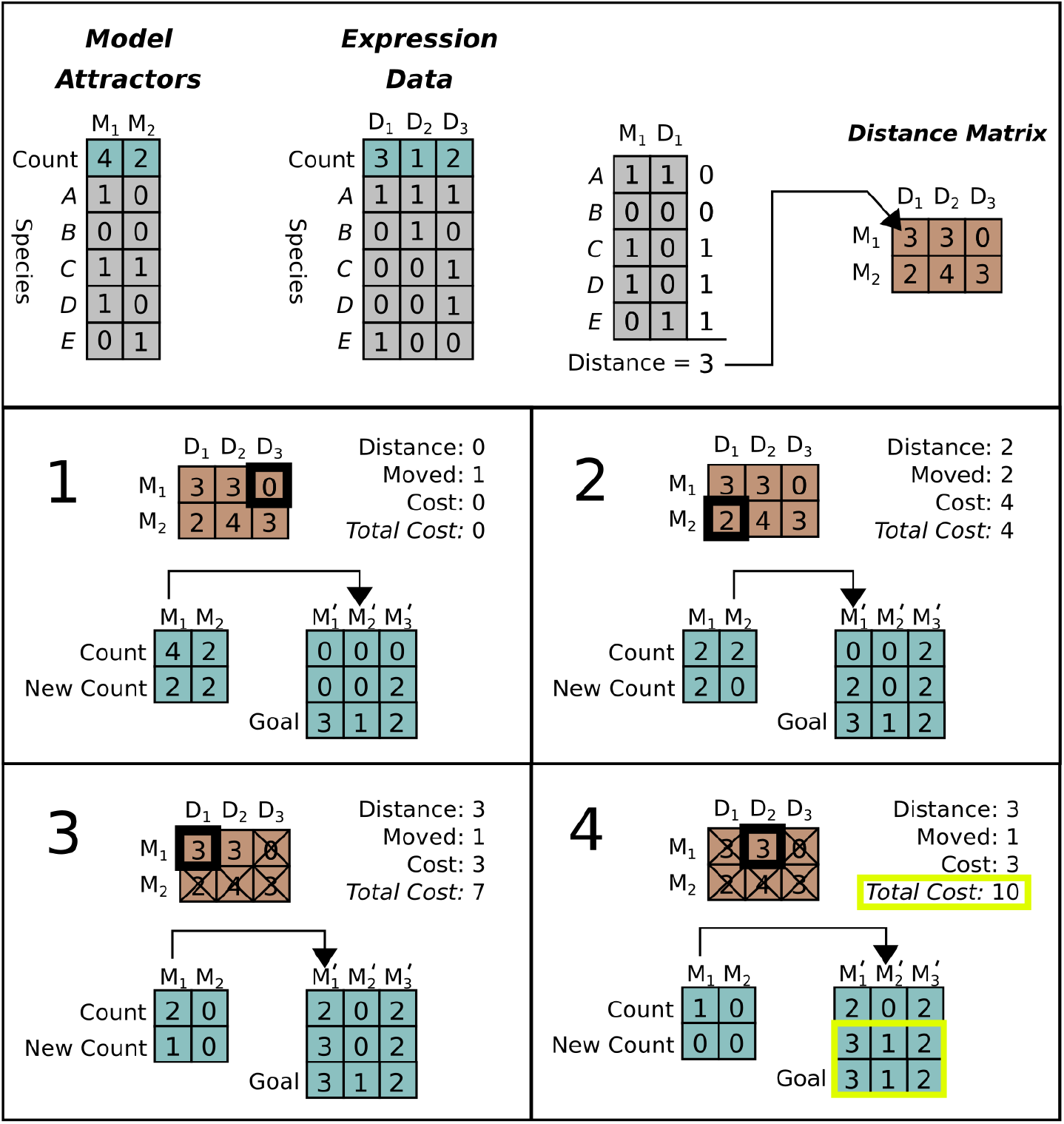
Example distance calculation for a system with five genes. *Top row:* Example attractors (left) are generated by simulating a model. The expression data (middle left) against which simulated attractors will be compared is also shown. Note that the data has three unique attractor states denoted *D*_*i*_ while the simulation only has two, denoted *M*_*i*_. To calculate the first entry in the distance matrix (right) attractor states *M*_1_ and reference states *D*_1_ are compared. Differences are assigned a “1” while matches are assigned a “0”. As shown, the distance between states *M*_1_ and *D*_1_ is “3” because they differ at three genes (C,D,E). *Bottom boxes:* Sequence of edits required to calculate the distance between simulation attractors (*M*) and data attractors (*D*). In the first step (1), we choose an edit by selecting the smallest valid distance from the distance matrix. This edit changes one of the *M*_1_ attractors to *D*_2_, but these are already identical, so the cost is zero. In step two (2) we select the next smallest distance (*M*_2_ to *D*_1_, with distance two) and change two attractors for a total cost of four. In step three (3) and four (4) we continue the same process. Note that in step three we remove multiple edits involving *M*_2_ from consideration, as all of the available *M*_2_ states have been edited already. In step four, the new state exactly equals *D*, so we halt the process with a final edit distance of ten.

At each step of the distance calculation, we identify every attractor state *s*_*i*_ in the simulation set where the occurrence count of the attractor is different between simulation and data (*c*_*i*_ ≠ *c*_*j*_). We then calculate “edits”, which is changing one attractor state to another. The cost *C* of an edit is the Manhattan distance between the bit vectors representing the state of the individual species in each attractor. We then find the edit with minimum cost that would maximally reduce *c*_*i*_ *− c*_*j*_. We apply these edits by changing the occurrence count of the edited simulation state, then repeat the process until all occurrence counts are equal between simulation and data. By accumulating edit costs at each step we obtain a total edit distance between simulated and measured attractor sets. This is normalized between (0, 1) by dividing by the maximum possible edit distance |*s*_*i*_ | *N*_*c*_, where |*s*_*i*_ | is the number species in the model and *N*_*c*_ is the sum of occurrence counts.

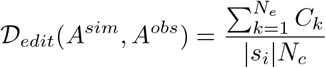

where *N*_*e*_ is the number of edits required, and *C*_*k*_ is the cost of the edit at step *k*. A detailed example of the edit distance calculation is shown in Fig 2.

### 3.3 Monte Carlo Tree Search

The core of MC-Boomer is Monte Carlo Tree Search (MCTS). In short, our MCTS procedure iteratively adds interactions to a Boolean model’s update rules, simulates the model, and then computes the similarity of simulated attractors to reference data. Each unique combination of rules is represented by a branch of the search tree. In Figure 3, each branch of the search tree is annotated with the unique set of interactions that comprise the corresponding model. On the left side of Figure 3 we show the Boolean models corresponding to the grey labeled nodes (M1-M3). MCTS probabilistically chooses which branches to continue expanding, based on a statistical upper bound on the similarity score of models from each branch. The upper bound is called the UCT or Upper Confidence bound for Trees. The upper bound is approximated by tracking the number of times a branch has been explored (visit count) and the average similarity scores of models on each branch of the search tree. These statistics and an example upper bound are shown for each node in the search tree in Figure 3.

**Figure 3:**
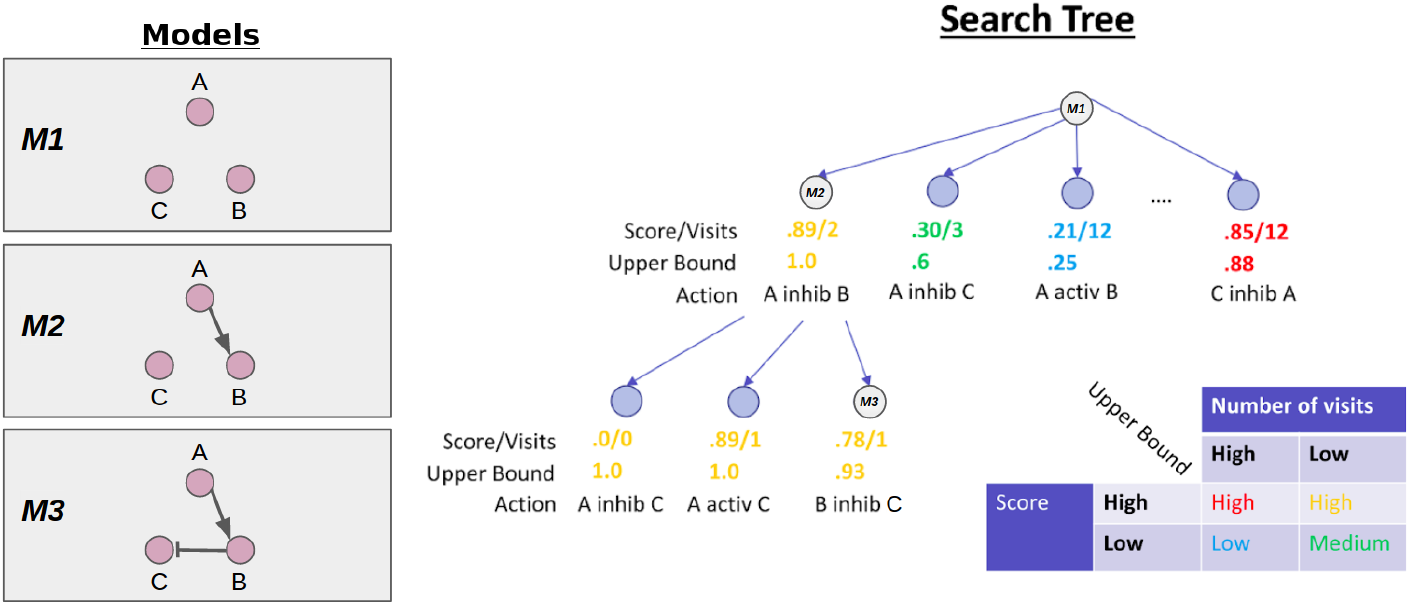
Overview of MCTS. On the left are the Boolean models corresponding to the branch of the search tree shown on the right, denoted M1, M2, M3. At each node in the tree, we also show the average score of models on the branch and the number of times the MCTS algorithm has visited the branch. These statistics are used to calculate the upper bound. In the bottom right, we show a conceptual overview of the functional form of the upper bound. In short, MCTS will aggressively explore branches with high scores but low number of visits. More exploration (i.e. a higher visit count) will progressively lower the upper bound until MCTS chooses another branch to explore.

MCTS uses the upper bound to balance exploration of different rules versus exploitation of rules that have already produced high scores. This is illustrated in Figure 3. The leftmost branch is relatively unexplored but models on that branch have high average similarity to the data. Thus, this branch has a high upper bound and the MCTS algorithm will preferentially explore and expand it. In contrast, the middle left branch has low average similarity scores so MCTS has “pruned” it from the search, leaving it unexplored. The middle right branch has low similarity, but has been explored several times, yielding a very low upper bound. Finally, the rightmost branch has high scores, but has been visited many times, and so the upper bound is close to the average score.

We implemented several modifications to standard MCTS that have been shown to improve the algorithm’s performance. RAVE is a simple modification to the MCTS algorithm that shares value estimates of actions across all branches of the search tree (Gelly and Silver, 2011). Nested search uses the actions from the best random rollout to choose the next step, rather than selecting based on upper confidence bound (Rosin, 2011). Branch retention keeps the upper confidence bound from previous search iterations and reuses them for every subsequent search step. These methods are described in more detail in the supplementary methods.

### 3.4 Validation Experiments

We performed two experiments to demonstrate MC-Boomer for inferring Boolean models. In sections 3.4.2 and 3.4.3, we randomly generated Boolean models of various sizes, then tested MC-Boomer’s ability to recover the structure and behavior of the random models. Then, in section 4.1.1, we tested MC-Boomer’s ability to recover the structure and behavior of the drosophila segment polarity network, a complex multicellular model that accurately recapitulates one aspect of drosophila embryo morphogenesis.

#### 3.4.1 Random Model Generation

Here we describe our validation experiments, showing that MC-Boomer can find models with a wide variety of behaviors and structures. We tested this by randomly generating models, simulating them, and then applying MC-Boomer to generate models matching their steady states.

We randomly generated Boolean models with dominant inhibition update rules by sampling uniformly from a list of all possible interactions between sets of 8 or 16 species. Following this procedure, we generated 80 random networks.

Before testing MC-Boomer on the randomly generated models, we ensured that the attractor states of the random models had realistic, diverse characteristics. The attractors reached by the random models do not collapse to an all active or inactive state, and instead have roughly one third active species (as shown in supplementary table S2). We consider the characteristics of these attractors to be biologically relevant, similar to data that might be obtained from an experiment. Thus, good performance on these random models shows that

MC-Boomer can generalize to a realistic variety of input data distributions.

#### 3.4.2 Steady State Behavioral Similarity

We applied MC-Boomer to attempt to recover these random models using only their initial states and attractors as input data. Figure 4a shows that the models generated by MC-Boomer at the beginning of the search process poorly matched the behavior of the ground truth models. This is expected, as the MCTS algorithm is effectively a random search process during the initial steps. However, by the end of the search, MC-Boomer reliably found models that had steady states with high similarity to the ground truth models. Across all model sizes, MC-Boomer was able to find several exact behavioral matches, with a majority having ¿95% similarity, as shown in Figure 4b.

**Figure 4:**
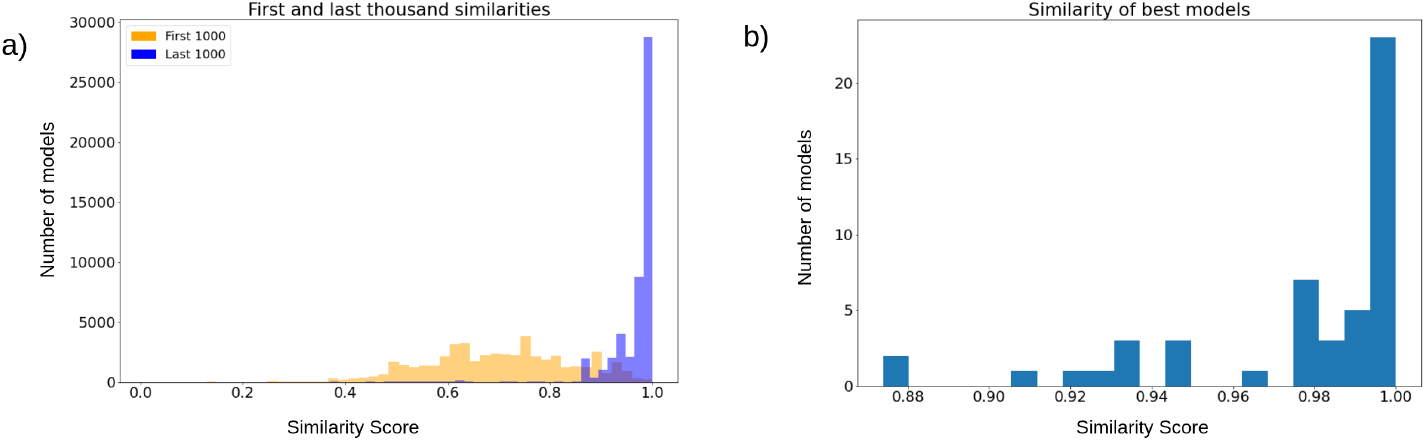
a) Orange histogram depicts distribution of similarities from the first one thousand models sampled during the MCTS search. Blue histogram is the distribution of similarities from the last thousand models. The blue distribution is significantly shifted towards higher rewards, indicating that MCTS was systematically sampling good models. b) Distribution of highest reward obtained by each independent search process. Most searches found models with ¿90% similarity.

#### 3.4.3 Rule Set Similarity

In addition to the steady state behavior of the models, we are also concerned with the content of the update rules generated by MC-Boomer. Many possible rule sets can have the same steady state behavior. However, many of these rule sets may be significantly different from each other and, most importantly, different from the underlying biological system. Under novel perturbations or conditions, these models may behave in radically different ways. Thus, we would like MC-Boomer to find models that match both the steady state behavior and the “interaction topology” of the underlying system. To validate MC-Boomer in this regard, we tested its ability to generate models with interactions that are similar to the reference models. In our tests, we quantified similarity by converting the update rules to sets of interactions for both the reference (randomly generated) model and the model generated by MC-Boomer. We then find the Jaccard index between the two interaction sets. This process is illustrated in Figure 5. Higher Jaccard indexes indicate that the MC-Boomer model matches the reference topology well.

**Figure 5:**
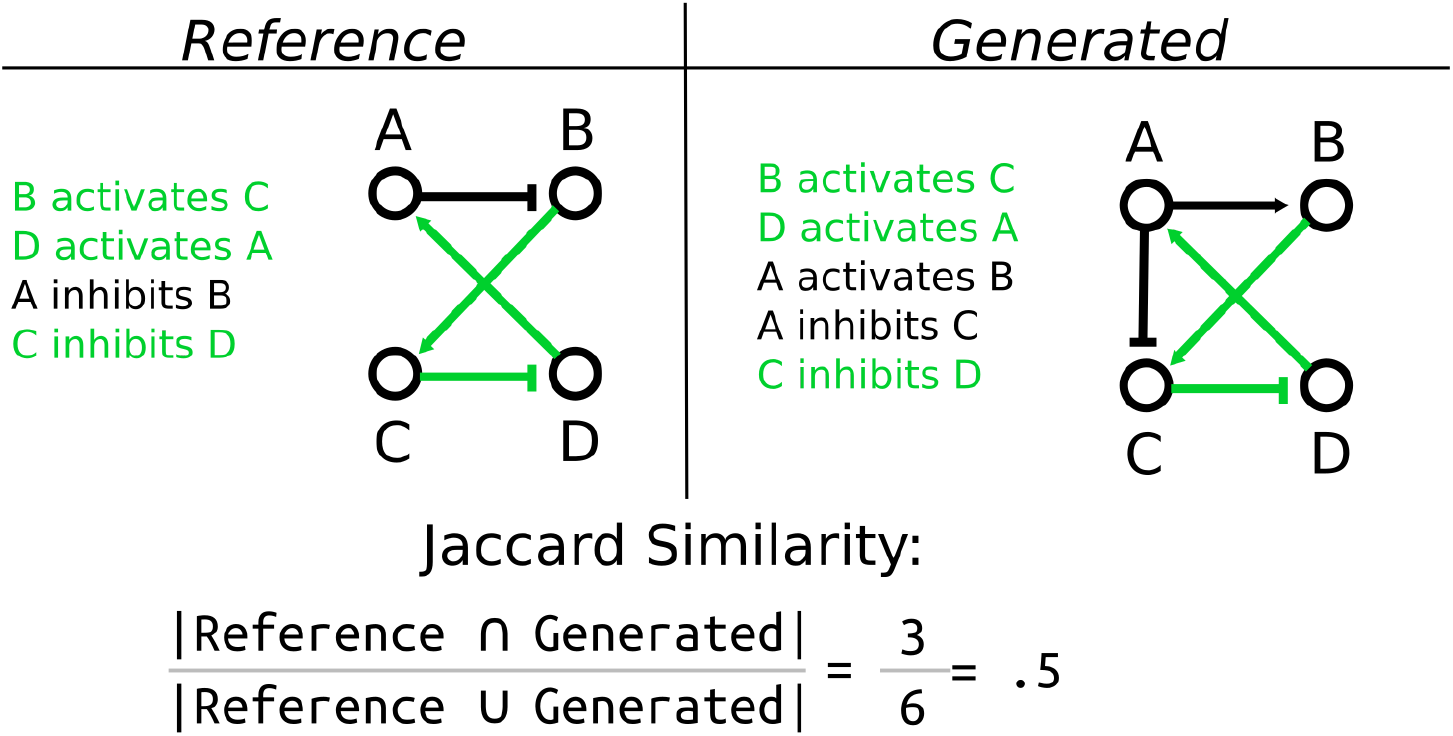
Example calculation of Jaccard similarity. We compare the structural similarity between two Boolean models by computing the Jaccard similarity between their sets of interactions. Here, shared interactions between the two models are highlighted in green, while interactions that are unique to each model are in black. In this example, the two models share three interactions in common, but have three more that are unique to each model. Thus they have a Jaccard similarity of 3*/*(3 + 3) = 3*/*6 = 0.5

With no restriction on the interactions selected by the model search process, MC-Boomer was able to find models with behavior that exactly matched the steady states of the reference models, but using rule sets that differed by as much as 80%. This corresponds to the left-most column of Figure 6, with zero reduction in search space, indicating that MC-Boomer was generating models using all possible interactions and no bias towards the true reference interactions.

**Figure 6:**
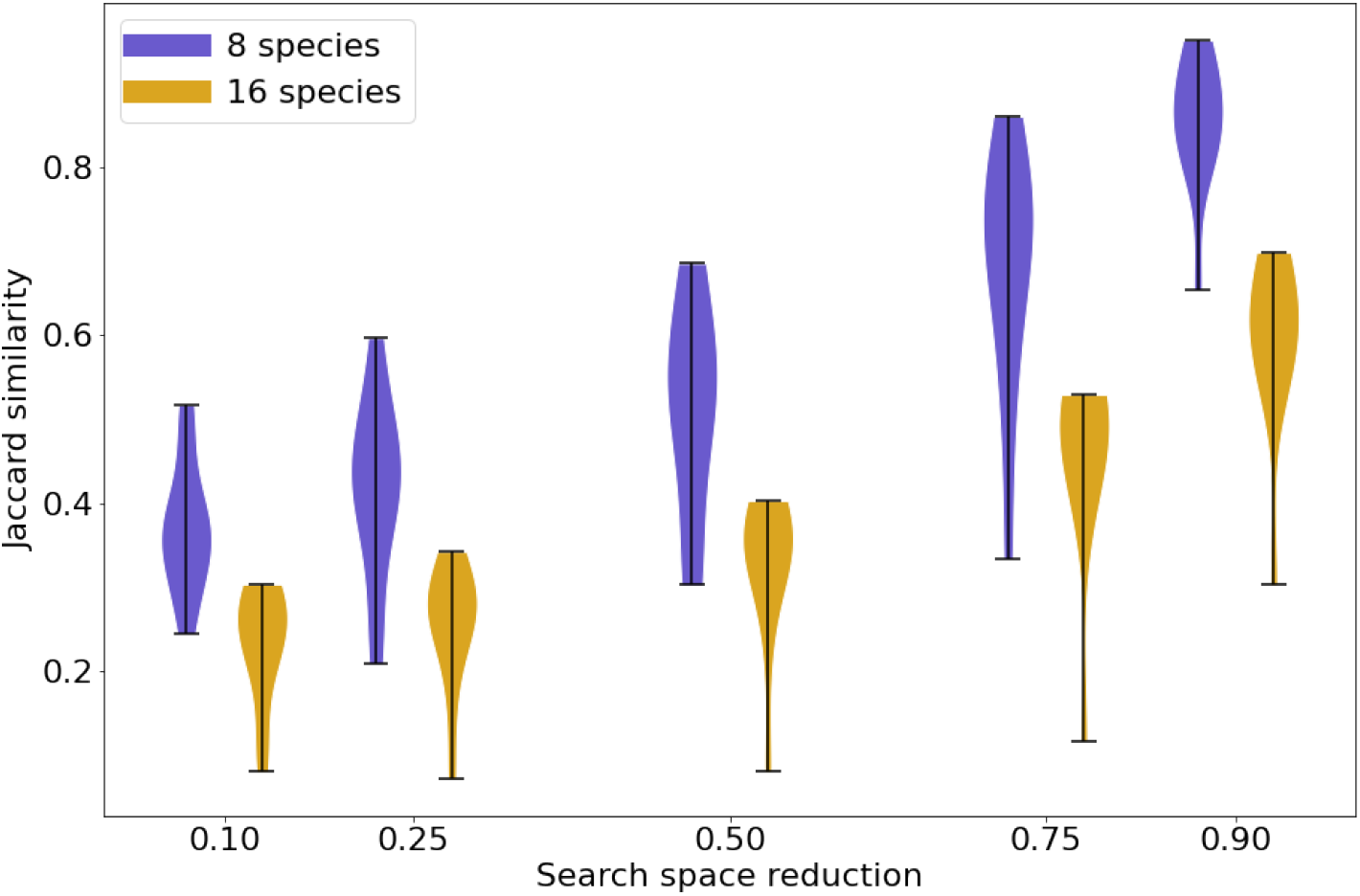
Jaccard similarity between synthetic reference and generated models with varying levels of prior knowledge. The violin plots show the distribution of Jaccard similarities achieved by MC-Boomer for synthetic models. The horizontal axis shows varying proportions of incorrect interactions randomly removed from the list of actions that MC-Boomer can choose when generating models. Removal of incorrect edges simulates the effect of prior knowledge, for example using only interactions from a database of validated biochemical interactions. As expected, higher levels of prior knowledge lead to higher Jaccard similarities, as MC-Boomer has a higher probability of choosing correct interactions from a smaller list.

We next investigated the effect of utilizing “prior knowledge” on MC-Boomer’s ability to recover correct rules. As noted above, model inference is an underconstrained problem with many possible models having data-consistent behavior, and so ruling out infeasible interactions can reduce the number of spurious models. We simulated varying levels of prior knowledge by randomly removing incorrect interactions from MC-Boomer’s action list, while retaining all of the correct interactions. We repeated the search five times, removing 10%, 25%, 50%, 75%, and then 90% of incorrect interactions from a set of 80 models. The aggregated Jaccard similarities for each percentage are shown in Figure 6. For models with both 8 and 16 species, increasing prior knowledge increased the Jaccard similarity to the reference data, as expected. Note that most proteinprotein interaction databases are much sparser than our highest tested level of prior knowledge. For example, BioGRID (version 4.4.2021) has 26k genes and 806k interactions, which corresponds to a 99.9% reduction from all possible interactions (Oughtred *et al*., 2021). Thus, our tests simulate a very difficult scenario, relying on much less prior knowledge than is available in biochemical interaction databases.

## 4 Results

Here we show the result of applying MC-Boomer to the segment polarity network (SPN). In sections 4.1.1 and 4.1.2, we describe the SPN and show MC-Boomer can generate models that are structurally similar to it, automatically discovering interactions that were previously manually selected by experts. Additionally we describe the large collection of alternate mechanisms generated by MC-Boomer, analyzing several in detail.

### 4.1 Segment Polarity Network (SPN)

As shown in the previous sections, MC-Boomer is able to generate models that are behaviorally and structurally similar to a variety of synthetically generated reference systems. While this was useful for validation, we also applied MC-Boomer to a more realistic setting to demonstrate the usefulness of the proposed framework. To that end, we employed MC-Boomer to build models of the *Drosophila* Segment Polarity Network (SPN), which is a gene circuit that controls the formation of borders and directionality of body segments during development of the drosophila embryo. As a reference, we have chosen a well-studied model by Albert and Othmer (Albert and Othmer, 2003). Briefly, this model comprises four cells, with several distinct components, including genes, proteins, membranes, protein isoforms, and complexes. A diagram of the SPN interactions is shown in Figure 7, and a complete listing of the reference rules are shown in supplementary table S3. Albert and Othmer provided binarized expression levels for wild type conditions as well as three gene knockouts, shown in supplementary table S4. We applied MC-Boomer with these expression profiles to automatically generate models of the SPN.

**Figure 7:**
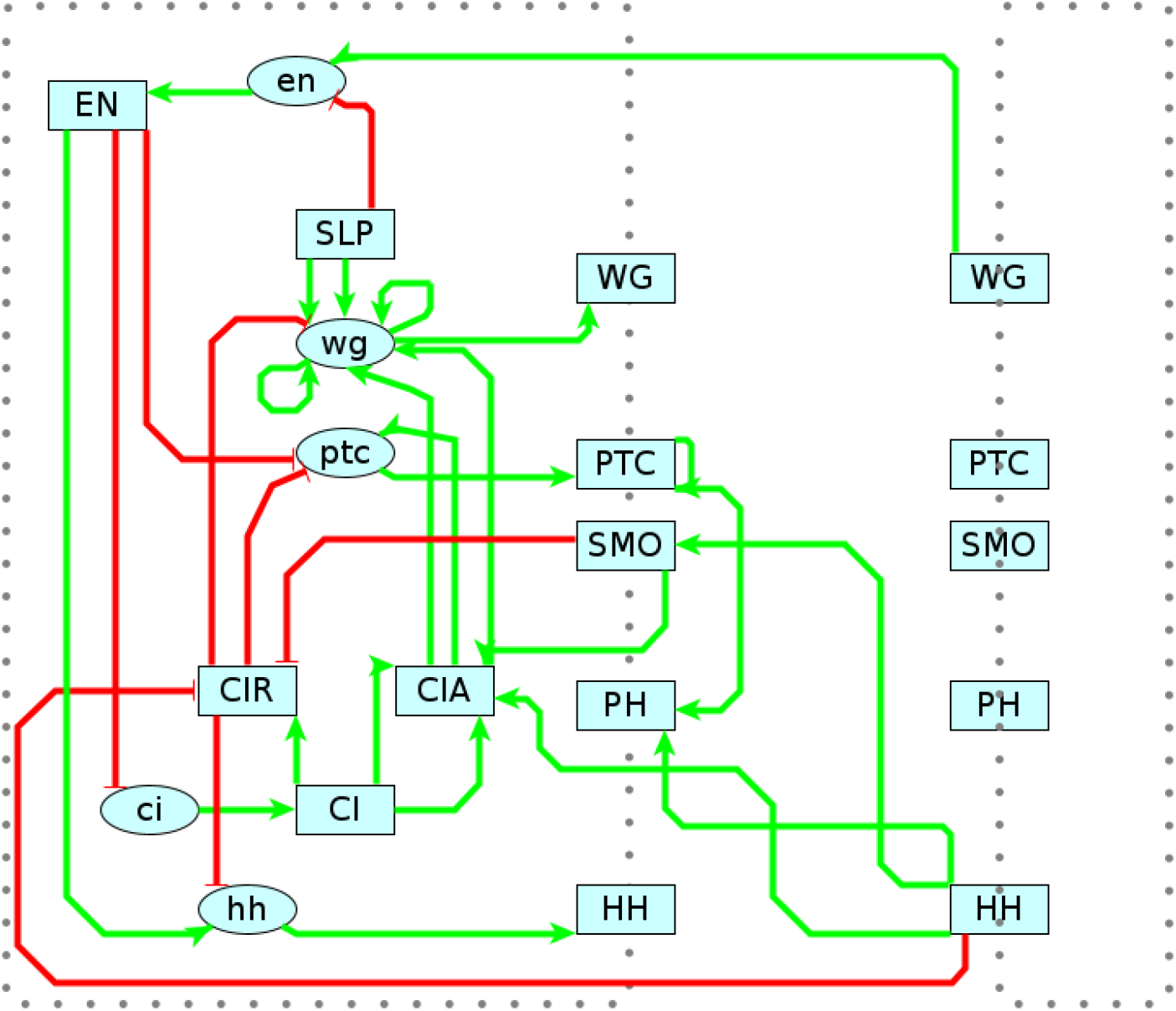
Reference Model for Segment Polarity Network. Diagram of the interactions in Albert and Othmer’s model of the segment polarity network (Albert and Othmer, 2003). Green edges indicate activating interactions. Red are inhibiting. Lower case ovals indicate genes and upper case indicate proteins. The dotted border indicates the cell membrane, with membrane proteins straddling the border. On the right is the adjacent cell, with several interactions spanning between cells. Albert and Othmer’s model has four cells with the same interactions inside each cell. Interactions between cells are symmetric, though only one direction is shown in the diagram to maintain clarity.

#### 4.1.1 Model Generation

First, we will describe how we initialized the model and performed the search. We applied several constraints to the search process so that MC-Boomer would only generate biologically plausible models. Membrane proteins (*WG, PTC, SMO, PH, HH*) could interact with membrane proteins only on adjacent cells. Internal proteins (*EN, SLP, CI, CIR, CIA*) could interact with other internal proteins, membrane proteins in the same cell, and genes in the same cell. Genes (*en, ci, ptc, hh*) could only activate their corresponding protein, and these gene-protein activating interactions were pre-specified in our search process.

All interactions added during the search process were repeated across all four cells. Multi-cellular membrane interactions were symmetric, added in both directions between neighboring cells.

The reference SPN model specified initial and stable states for the wild type network as well as initial and attractor states for knockouts of *wg, hh*, and *en* (see Supplementary figure S4 for details).

We applied MC-Boomer to search for models that matched the behavior of the reference SPN model across the wild-type and three knockout conditions. At each search iteration, we simulated the model across all conditions, calculated the edit distance between simulated and reference steady states, then averaged all conditions’ similarity scores to get a final score for the iteration. We implemented knockouts by removing all interactions to and from the *hh, en* and *wg* genes across all cells.

We ran 1500 searches in batches of 30 in parallel on our institution’s computing cluster. In each search step, MC-Boomer simulated 10k model variations before adding the best interaction to the model and starting the next step. We restricted the search to terminate after 30 steps, but not before completing 8 steps. Every search was run with RAVE, nested search, and branch retention enabled with the same uniformly random sampled parameter distributions as in the synthetic data experiments. The complete search process took 41 hours and simulated 430 million unique models. Eleven of the 1500 search processes found models with exactly the same steady states as the reference model for all four conditions. Collectively, these eleven search processes generated ¿202*k* models with perfect consistency to the attractor data.

#### 4.1.2 Visualizing the Set of Data-Consistent Models

Given the our collection of models with consistent steady state behavior, we were motivated to develop methods for visualization and exploration of large numbers of models.

First, we applied dimensionality reduction and clustering methods to visualize similarities between the models. We randomly sampled fifty thousand of the 202*k* data-consistent models and clustered them with the UMAP algorithm (McInnes *et al*., 2018) using the interaction set Jaccard distance between models, as illustrated in Figure 5. Model sampling was necessary because UMAP requires computation of a pairwise distance matrix that would have been infeasible for the full data set. Multiple different samples all gave similar results, thus providing us with confidence that the sample analyzed here was representative of the overall model population.

Applying UMAP with the Jaccard distance yielded the result shown in Figure 8 with eleven well separated clusters, corresponding to the eleven independent searches that produced data-consistent models.

**Figure 8:**
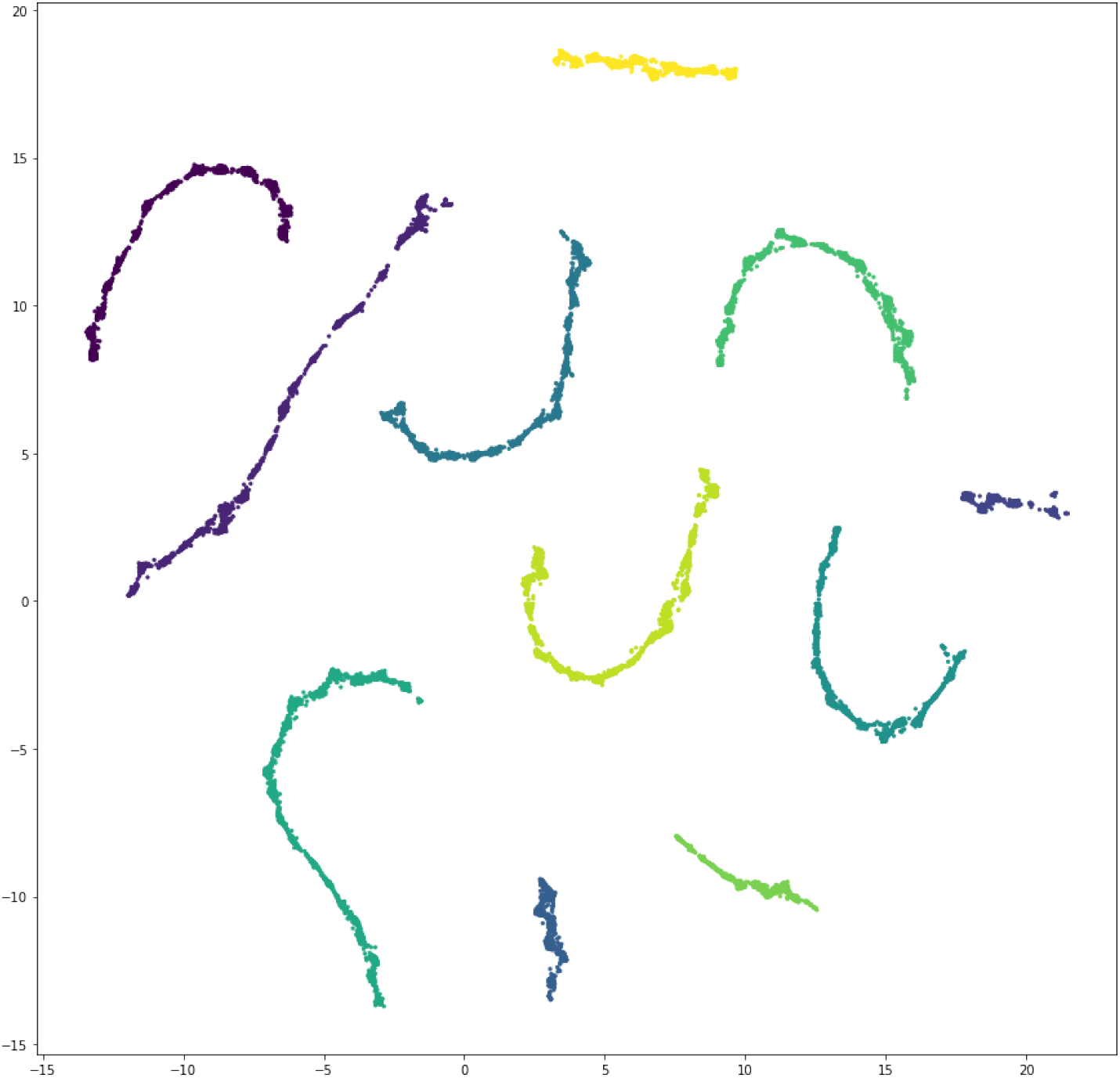
Scatter plot depicting clustering of unique data-consistent segment polarity models after UMAP projection to two dimensions. There are eleven well separated clusters, corresponding to the eleven independent search processes that found data-consistent models.

**Figure 9:**
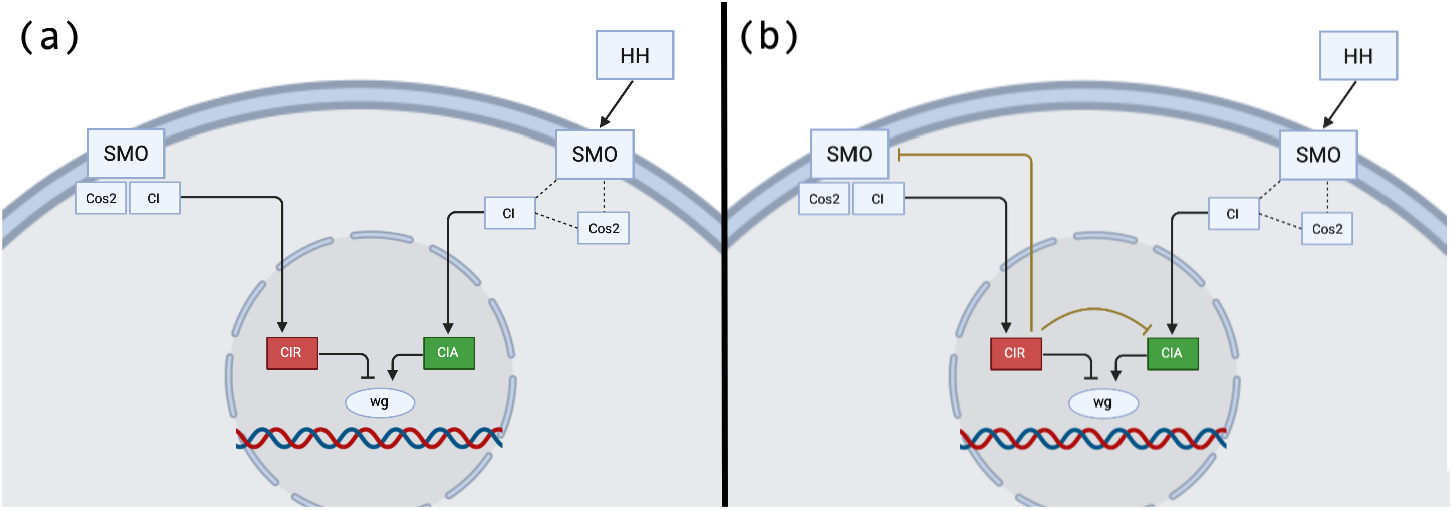
a) The reference model depicts the modification of CI as a forked pathway, where the resulting product is determined by the activation state of SMO. In the reference model, active SMO promotes CIA and inhibits CIR. b) The MC-Boomer model, in contrast, includes two novel interactions where CIR inhibits CIA and SMO. Figures partially based on Figure 3 from Hooper and Scott (2005)

**Table 1:**
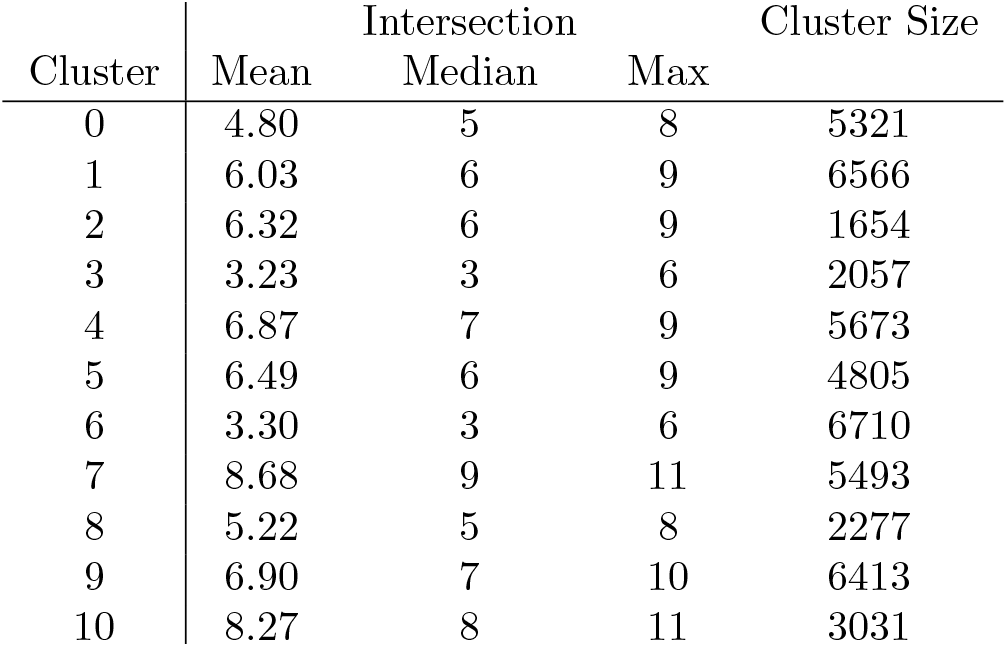
Structural Intersection with Reference Model

Table 2: For each cluster of models shown in Fig. 8 we computed the intersection between these common rules and the reference model. We show the mean, median, and maximum intersection between each cluster’s models and the reference. Cluster 7 has the highest intersection across all three statistics, while cluster 3 shares the fewest interactions with the reference. We further investigate the most common interactions in Cluster 7 in Fig. 10.

#### 4.1.3 Structural Similarity between Clusters and Reference

We then compared the interactions in each MC-Boomer generated model with the interactions in the reference model’s update rules to find the “structural” similarity.

**Figure 10:**
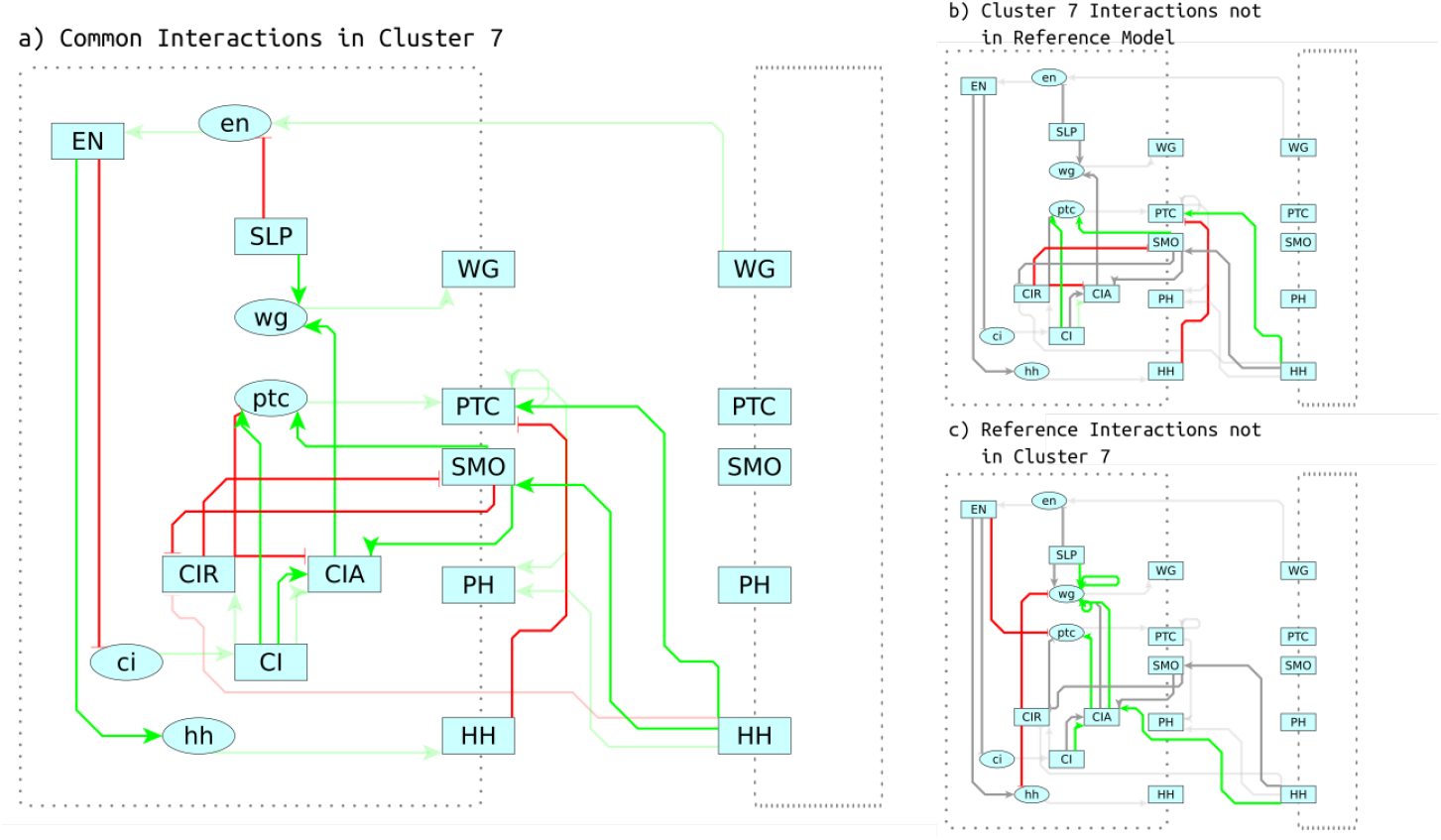
a) Most common interactions in cluster 7. These interactions that are found in ¿90% of models in cluster 7. Transparent lines represent interactions that were pre-specified as prior knowledge, while more opaque interactions were generated by MC-Boomer. b) Interactions highlighted in red and green are common in cluster 7 but are not present in the reference model. Grey interactions are shared between the reference model and cluster 7. Note the red inhibiting interaction between CIR and CIA, which is investigated in more detail in Section 4.1.2 c) The reference model is shown here, highlighting interactions that are in not present in cluster 7. Again, note the red inhibiting interaction between CIR and wg. Section 4.1.2 proposes an alternative mechanism for inhibition of wg by CIR.

The update rules of the reference model had 26 total interactions. We manually pre-specified eleven of the interactions in the reference segment polarity network. That is, all models generated by MC-Boomer included these interactions as “prior knowledge”. This included all the interactions in which a gene activated its corresponding protein, as well as four interactions that did not fit the dominant inhibition dynamics of the rest of the network (supplementary figure S1). Our tests evaluated MC-Boomer’s ability to discover models that included the remaining 15 interactions in the reference model.

Within each cluster of MC-Boomer models, we computed the mean, median, and maximum size of the intersection between the cluster’s models’ interactions and the reference model’s interactions, as shown in Table 2.

Comparison across clusters revealed a wide disparity in accuracy, with cluster 3 having, on average, 3 rules in common with the reference SPN model. We note that while the models in cluster 3 had low structural similarity to the reference SPN model, all of the models in every cluster have the same steady state attractors as the reference. Cluster 7 had the highest average intersection, with several models in the cluster having 11 out of 15 rules in common with reference model. For cluster 7, we found the most common rules, i.e. those shared by ¿90% of the models in the cluster. Figure 10a shows these common rules and Figure 10b/c shows “false positive” and “false negative” rules, respectively. False positive were present in MC-Boomer models but not in the reference and false negative rules were in the reference but not the MC-Boomer models. In the following sections, we investigate two of these interactions, one of which was not present in the reference model.

#### 4.1.4 Investigating Common Mechanisms

The high-level clustering analysis shows that MC-Boomer generates models with a wide variety of structures but identical steady state behavior. However, this analysis is too broad to elucidate the precise nature of the mechanisms that these models employ to generate this behavior. Accordingly, we more closely investigated two key interactions that are present in every model generated by MC-Boomer. Specifically, we consider “*EN* inhibits *ci*” and “*CIR* inhibits *CIA*”, which are present in 100% of the data-consistent models.

First, we look at *EN* inhibiting *CI*, which is present in all of our models and also present in the reference model. This indicates that this interaction is a crucial link across the the very diverse mechanisms employed by the eleven clusters of models and the reference model. Simulating a random sample of one thousand models with this interaction knocked out resulted in a 28% average absolute reduction in similarity to the reference steady state data. We observed that knocking out the *EN* to *ci* interaction in the reference model also reduced similarity to the reference data by 28%. Again, this indicates that the while the models are structurally diverse, they share a similar reliance on this particular interaction of *EN* and *ci*.

On the contrary, CIR inhibition of CIA is not present in the reference model. This interaction is shared by more than two hundred thousand unique models generated by MC-Boomer. The high frequency of the CIR inhibiting CIA interaction motivated further investigation into CIR and CIA’s role in regulation of the wg gene.

To provide necessary background for our discussion of wg regulation, we briefly describe the key genes in this pathway. CIA is an activated, nuclear transported form of the CI protein, while CIR is a proteolytically cleaved form of CI which represses wg transcription. In the absence of HH, SMO forms a complex with CIA and Cos2, a kinesin-like protein that binds and sequesters CIA, preventing its nuclear translocation and permitting its cleavage into CIR. In the presence of HH, SMO is activated and Cos2 releases CIA, which is then transported to the nucleus, where it activates wg (Lum *et al*., 2003; Kalderon, 2004; Ranieri *et al*., 2012). The exact mechanisms and network dynamics behind CI activation, cleavage, and nuclear translocation have long remained a point of debate and uncertainty (Ruel *et al*., 2003).

In addition to CIR inhibiting CIA, MC-Boomer also suggests (in 34% of models) an inhibitory interaction between CIR and SMO. The novel inhibition of CIA and SMO by CIR can be interpreted in at least two ways.

1. These interactions do not represent real signaling mechanisms. In accordance with the reference model, the bi-directional inhibitory loop between CIR and SMO may simply reflect the normal activation states of these proteins. When SMO is active, CIR cannot be produced because SMO destabilizes Cos2 and therefore all CI is available as CIA. Conversely, when SMO is inactive, Cos2 binds CI and conversion to CIR occurs. Therefore, the inhibition of CIA and SMO by CIR may not represent genuine biochemical interactions, but may simply be artifacts of MC-Boomer’s automated model generation process.
2. These interactions do represent real, redundant signaling mechanisms. The novel inhibition of CIA and SMO by CIR may represent redundant signals which prevent the possibility of competition at the target gene binding site. This type of redundancy is a feature observed in other biological signaling networks (Albert 2011). CIR inhibition of CIA and SMO in the cytosol ensures that CIR can bind and inhibit wg in the nucleus without interference from CIA. In this interpretation, CIR is not just a passive cleavage product, but also an active participant in a feedback loop that inhibits the activity of CIA.

This second interpretation describes an instance of signaling redundancy. If CIR inhibits SMO and CIA, this helps to ensure a full transition between on and off network states and prevents any potential binding competition at the target gene.

Overall, these observations show that the proposed method can both reproduce the known biological features as well as provide novel insight into the segment polarity network by generating new mechanistic hypotheses, which require further investigation through experiments.

#### 4.1.5 Identifying Unique Mechanisms in Model Clusters

We are able to analyze these two interactions in detail because they are shared across all models and their limited scope eases their interpretation. However, our clustering analysis showed that there are at least eleven groups of models with widely differing structures. Accordingly, we also investigated the role of interactions that are specific to individual clusters of models. We searched for sets of up to 5 interactions that are present in a high proportion of models in each cluster, while not being present in models in other clusters. We call these distinguishing sets. We found between 25 and 21,570 distinguishing sets per cluster.

Given the large number of distinguishing sets for some clusters, we needed a measure of which sets are most important to the function of the models in the cluster. We quantified this by simulating knock outs of each distinguishing set in a sample of 100 models from their respective clusters and calculating the reduction in similarity to the reference data caused by the knockouts. We refer to distinguishing sets with the largest reduction in similarity as the maximally disruptive sets. These maximally disruptive sets identify the unique mechanisms that the models in each cluster most highly rely on to generate their behavior. Comparing the interactions in the maximally disruptive sets revealed heterogeneity across the clusters. Most of the maximally disruptive sets shared two or fewer interactions in common. For example, the maximally disrupting sets for cluster 7 (Figure 11a) and cluster 8 (Figure 11c) only share a single interaction in common. Simulated knockouts of cluster 7 and 8’s maximally disruptive sets reduced similarity to reference data by 31% and 39%, respectively. This indicates that models in these two clusters depend, to a similar degree, on these distinct sets of interactions for generating correct behavior. Inspection reveals that while the two mechanisms are not similar by a direct comparison, they share functional similarity in primarily modulating the connectivity and activity of *EN*. This corresponds with our previous analysis showing that *EN* interactions are crucial for correct model behavior across our whole collection of models. However, the actual mechanism by which *EN* activity is directed is quite distinct. The interactions in cluster 7 (Figure 11a) give *EN* a mixed activating/inhibiting role, while cluster 8 (Figure 11b) relies on several inhibitory feedback loops centered on *EN*.

**Figure 11:**
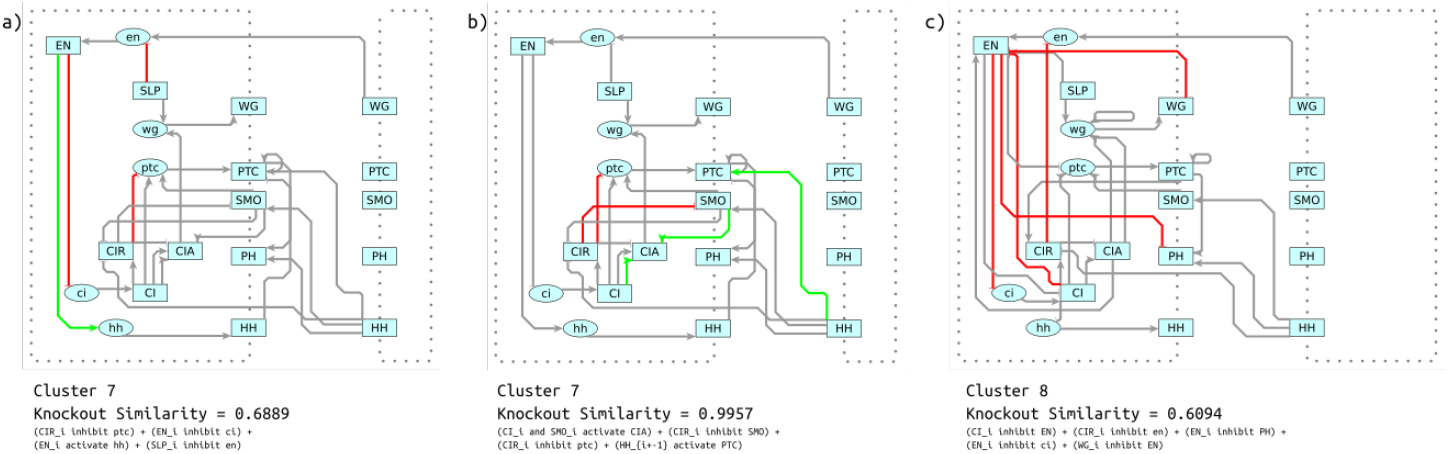
a) Shown in red and green is the maximally disruptive set for cluster 7. These are interactions that are common in cluster 7, but are very uncommon in other clusters. Additionally, knocking out these interactions reduces similarity to the reference steady states more than other sets of common interactions (shown in grey). b) Another distinguishing set of interactions, but these are minimally disruptive. Knocking them out only reduces similarity to the reference data by a negligible amount. c) Maximally disruptive set for cluster These reduce similarity to reference data to a similar degree as the most disruptive set in cluster 7, but these interactions utilize a different mechanism.

Similar to the case of CIR described in Section 4.1.4, many of the distinguishing sets do not have any effect on the behavior of the model; one such example is illustrated by Figure 11b. One perspective is that these interactions are redundant and only increase the complexity of the model. Accordingly, several previous approaches (citations) penalize models with more interactions. Another perspective is that these redundant connections may confer robustness, i.e. an ability to recover from aberrant initial conditions or losses of function, or as with CIR they could help ensure full response to inhibition or activation.

## 5 Discussion

Biology is inherently complex, yet our measurements capture only a limited slice of the true activity within a cell. Current assay technology can only describe a subset of biomolecules at low time resolution and with significant noise. From this blurry view researchers must synthesize a model that can both describe the phenomena under investigation and predict the system’s behavior in novel circumstances. Synthesizing a model can be made easier by choosing the simplicity of the Boolean logic modeling formalism to represent the system. Nonetheless, even for a small number of interacting species, the number of possible Boolean models is vast. Consequently, a typical researcher, creating models through trial and error, may only find one or perhaps a few models whose behavior is consistent with the observed data. However, as we have shown in Section 4.1.1, even in a small system with multiple measurements and reasonable prior assumptions on model structure, there are hundreds of thousands of models that are all consistent with the data.

This observation was made possible by using an efficient search technique, Monte Carlo Tree Search, to build models. We demonstrate the power of MCTS to synthesize models with the correct steady-state behavior and the correct interactions in Section 3.4. While previous studies have shown that similar optimization methods (e.g. tabu search in Aghamiri and Delaplace (2020)) are effective for finding data-consistent models, they have focused on finding a single model that is “best” in terms of both complexity and fit to the data. In contrast, we retain every model that fits the data well and in Sections 4.1.4 and 4.1.5 we develop a set of techniques for making sense of this large collection of models.

We approach this from a data-driven perspective, in the sense that our MCTS algorithm generates data about the space of valid hypotheses. By clustering models based on their structural features, we can find recurrent motifs across the whole collection of models, as well as distinct motifs that discriminate the structure of groups of models. Simulated knockouts of these motifs then reveal that some are critical to the models’ correct behavior.

### 5.1 Using MC-Boomer to Design Experiments

As we describe in Section 4.1.4, analysis of the models generated by MC-Boomer pointed us towards an alternate hypothesis for the mechanism by which *CIR* and *CIA* regulate expression of the wingless gene (*wg*) in the segment polarity network. An investigator using MC-Boomer to study this pathway may propose that CIA activation of *wg* depends on both SMO (Smoothened) stabilization and, as MC-Boomer suggests, the absence of CIR. The existence of these novel inhibitory relationships could be experimentally validated by introducing CIR into cells in which HH signaling has already activated SMO and CIA. Reduced concentrations of active CIA or SMO would indicate that CIR does, in fact, inhibit the activity of CIA and SMO.

### 5.2 Limitations and Future Work

Previous work (Fauré *et al*., 2006) has suggested that the general asynchronous updating scheme yields more biologically realistic results for Boolean network simulations. While our current approach uses synchronous updating, extending MC-Boomer to work with asynchronous updating would be straightforward.

The current approach is limited in its scalability to models with large numbers of interacting species by several key bottlenecks. First, this approach requires simulation of every synthesized model, and simulation becomes prohibitively expensive for large models. This could be alleviated through partial or approximate simulations of the models. While this would yield an approximation of the model’s similarity to data, the UCT upper bound allows MCTS to tolerate some noise in the search process. Second, the search space scales exponentially with the number of species in the model. We show that restricting the search space through prior knowledge constraints on model structure is an effective strategy for improving structural and behavioral accuracy of synthesized models. The efficiency of the search algorithm could further be improved by using deep learning to guide MCTS. This is similar to the approach used by the AlphaZero algorithm (Silver *et al*., 2018), that proved to be exceptionally effective at searching the combinatorially large space of moves in games like chess and Go. We are currently exploring each of research directions as potential optimizations of the MC-Boomer algorithm.

