## Supplementary Tables and Figures for "Automatic mechanistic inference from large families of Boolean models generated by Monte Carlo Tree Search"

### Supplementary Methods

#### MCTS Enhancements

##### Rapid Action Value Estimation (RAVE)

In standard MCTS, the estimated best value of a given node (i.e. adding a specific interaction to the Boolean model) is computed by backpropagating similarity scores from all random rollouts below that node in the search tree. Thus, the node's value estimate is limited to rollouts from its branch of the tree. However, other branches of the search tree may include the same action. Rollouts from the other branches could provide some amount of information about the value of the action on other branches. This is the motivating insight of RAVE. RAVE maintains a list of actions with their number of visits and best values, accumulated across all branches. During selection, a node's value is calculated as a weighted average of the RAVE value and the standard (branch specific) value.

$$\hat{\mathcal{D}} = (1 - \beta(n_i, k))\mathcal{D}^* + \beta(n_i, k)\mathcal{D}^{rave}$$

Here, $D^*$ is the branch-specific value, $D^{rave}$ is the accumulated RAVE value, and $\hat{D}$ is the weighted average. The weighting factor β is a heuristically determined function of the number of visits to search tree node $n_i$ and a parameter k that determines the number of visits when $D^*$ and $D^{rave}$ are weighted equally:

$$\beta(n_i, k) = \sqrt{\frac{k}{3N_v(n_i) + k}}$$

The weighted average is then substituted into the UCB calculation during selection:

$$\mathrm{UCB} = \hat{\mathcal{D}} + c\sqrt{\frac{\ln N_v(n_i)}{N_v(n_j)}}$$

##### Nested Search

In standard MCTS, after reaching the iteration limit, we add the best interaction to our model and restart the search process. As a result, information from the previous search steps is lost. We remedy this by retaining the sequence of actions taken by the rollout with the highest reward. If any subsequent search steps fail to find a rollout with higher similarity, then we take the next action from the previous best rollout sequence. This technique is known as nested MCTS.

##### Branch Retention

As explained above, nested search allows MCTS to retain information from previous search steps about the best sequence of actions. However, the statistics stored at each node of the tree are lost at each step. We can retain the search statistics for the branch chosen at each step, an option we refer to as "branch retention".

**Supplementary Table S1 - Number of attractors in randomly sampled models**

| | Stable | | Cyclic | | |
|---|---|---|---|---|---|
| | Mean±Std | Median | Mean±Std | Median | Cycle length |
| 8 species | 2.6±1.11 | 3.00 | 2.90±1.75 | 3.00 | 2.65±0.91 |
| 16 species | 2.4±1.30 | 2.00 | 3.59±2.47 | 3.00 | 3.37±1.91 |
| 32 species | 2.4±1.39 | 2.00 | 4.42±2.72 | 4.00 | 5.36±6.83 |

**Supplementary Table S2 - Active/Inactive ratio of attractors from sampled models**

| | Stable | | Cyclic | |
|---|---|---|---|---|
| | Mean±Std | Median | Mean±Std | Median |
| 8 species | 0.3±0.27 | 0.33 | 0.33±0.20 | 0.30 |
| 16 species | 0.2±0.21 | 0.19 | 0.28±0.17 | 0.30 |
| 32 species | 0.2±0.20 | 0.15 | 0.30±0.15 | 0.30 |

**Supplementary Table S3 - Segment Polarity Network Reference Rules**

$$
\begin{aligned}
SLP_i^{t+1} &:= \begin{cases} 0 & \text{if i} \in \{0,2\} \\ 1 & \text{if i} \in \{1,3\} \end{cases} \\
wg_i^{t+1} &:= (\text{CIA}_i^t \text{ and } \text{SLP}_i^t \text{ and not } \text{CIR}_i^t) \text{ or } [\text{wg}_i^t \text{ and } (\text{CIA}_i^t \text{ or } \text{SLP}_i^t) \text{ and not } \text{CIR}_i^t] \\
&= (\text{CIA}_i^t \text{ and } \text{SLP}_i^t \text{ or } \text{wg}_i^t \text{ and } \text{CIA}_i^t \text{ or } \text{wg}_i^t \text{ and } \text{SLP}_i^t) \text{ and not } \text{CIR}_i^t \\
WG_i^{t+1} &:= \text{wg}_i^t \\
en_i^{t+1} &:= (\text{WG}_{i-1}^t \text{ or } \text{WG}_{i+1}^t) \text{ and not } \text{SLP}_i^t \\
EN_i^{t+1} &:= \text{en}_i^t \\
hh_i^{t+1} &:= \text{EN}_i^t \text{ and not } \text{CIR}_i^t \\
HH_i^{t+1} &:= \text{hh}_i^t \\
ptc_i^{t+1} &:= \text{CIA}_i^{t+1} \text{ and not } \text{EN}_i^t \text{ and not } \text{CIR}_i^t \\
&= \text{CIA}_i^{t+1} \text{ and not } (\text{EN}_i^t \text{ or } \text{CIR}_i^t) \\
PTC_i^{t+1} &:= \text{ptc}_i^t \text{ or } (\text{PTC}_i^t \text{ and not } \text{HH}_{i\pm1}) \\
PH_i^{t+1} &:= \text{PTC}_i^t \text{ and } (\text{HH}_{i-1}^t \text{ or } \text{HH}_{i+1}^t) \\
SMO_i^{t+1} &:= \text{not } \text{PTC}_i^t \text{ or } \text{HH}_{i-1}^t \text{ or } \text{HH}_{i+1}^t \\
ci_i^{t+1} &:= \text{not } \text{EN}_i^t \\
CI_i^{t+1} &:= \text{ci}_i^t \\
CIA_i^{t+1} &:= \text{CI}_i^t \text{ and } (\text{SMO}_i^t \text{ or } \text{hh}_{i-1}^t \text{ or } \text{hh}_{i+1}^t) \\
CIR_i^{t+1} &:= \text{CI}_i^t \text{ and not } (\text{SMO}_i^t \text{ or } \text{hh}_{i-1}^t \text{ or } \text{hh}_{i+1}^t)
\end{aligned}
$$

#### Supplementary Table S4 - Wild Type and Knockout Initial and Attractor States

##### Wild type attractors

| | | | | |
|---|---|---|---|---|
| wg | 0 | 1 | 0 | 0 |
| WG | 0 | 1 | 0 | 0 |
| en | 0 | 0 | 1 | 0 |
| EN | 0 | 0 | 1 | 0 |
| hh | 0 | 0 | 1 | 0 |
| HH | 0 | 0 | 1 | 0 |
| ptc | 0 | 1 | 0 | 1 |
| PTC | 1 | 1 | 0 | 1 |
| PH | 0 | 1 | 0 | 1 |
| SMO | 0 | 1 | 1 | 1 |
| ci | 1 | 1 | 0 | 1 |
| CI | 1 | 1 | 0 | 1 |
| CIA | 0 | 1 | 0 | 1 |
| CIR | 1 | 0 | 0 | 0 |
| SLP | 1 | 1 | 0 | 0 |

##### Wild Type Initial State

| | | | | |
|---|---|---|---|---|
| wg | 0 | 1 | 0 | 0 |
| WG | 0 | 0 | 0 | 0 |
| en | 0 | 0 | 1 | 0 |
| EN | 0 | 0 | 0 | 0 |
| hh | 0 | 0 | 1 | 0 |
| HH | 0 | 0 | 0 | 0 |
| ptc | 1 | 1 | 0 | 1 |
| PTC | 0 | 0 | 0 | 0 |
| PH | 0 | 0 | 0 | 0 |
| SMO | 0 | 0 | 0 | 0 |
| ci | 1 | 1 | 0 | 1 |
| CI | 0 | 0 | 0 | 0 |
| CIA | 0 | 0 | 0 | 0 |
| CIR | 0 | 0 | 0 | 0 |
| SLP | 1 | 1 | 0 | 0 |

**Knockout Experiment Attractor (en, hh, wg knockout)**

| | | | | |
|---|---|---|---|---|
| wg | 0 | 0 | 0 | 0 |
| WG | 0 | 0 | 0 | 0 |
| en | 0 | 0 | 0 | 0 |
| EN | 0 | 0 | 0 | 0 |
| hh | 0 | 0 | 0 | 0 |
| HH | 0 | 0 | 0 | 0 |
| ptc | 0 | 0 | 0 | 0 |
| PTC | 1 | 1 | 1 | 1 |
| PH | 0 | 0 | 0 | 0 |
| SMO | 0 | 0 | 0 | 0 |
| ci | 1 | 1 | 1 | 1 |
| CI | 1 | 1 | 1 | 1 |
| CIA | 0 | 0 | 0 | 0 |
| CIR | 1 | 1 | 1 | 1 |
| SLP | 1 | 1 | 0 | 0 |

**Supplementary Figure S1 - Base model used in MC-Boomer search**

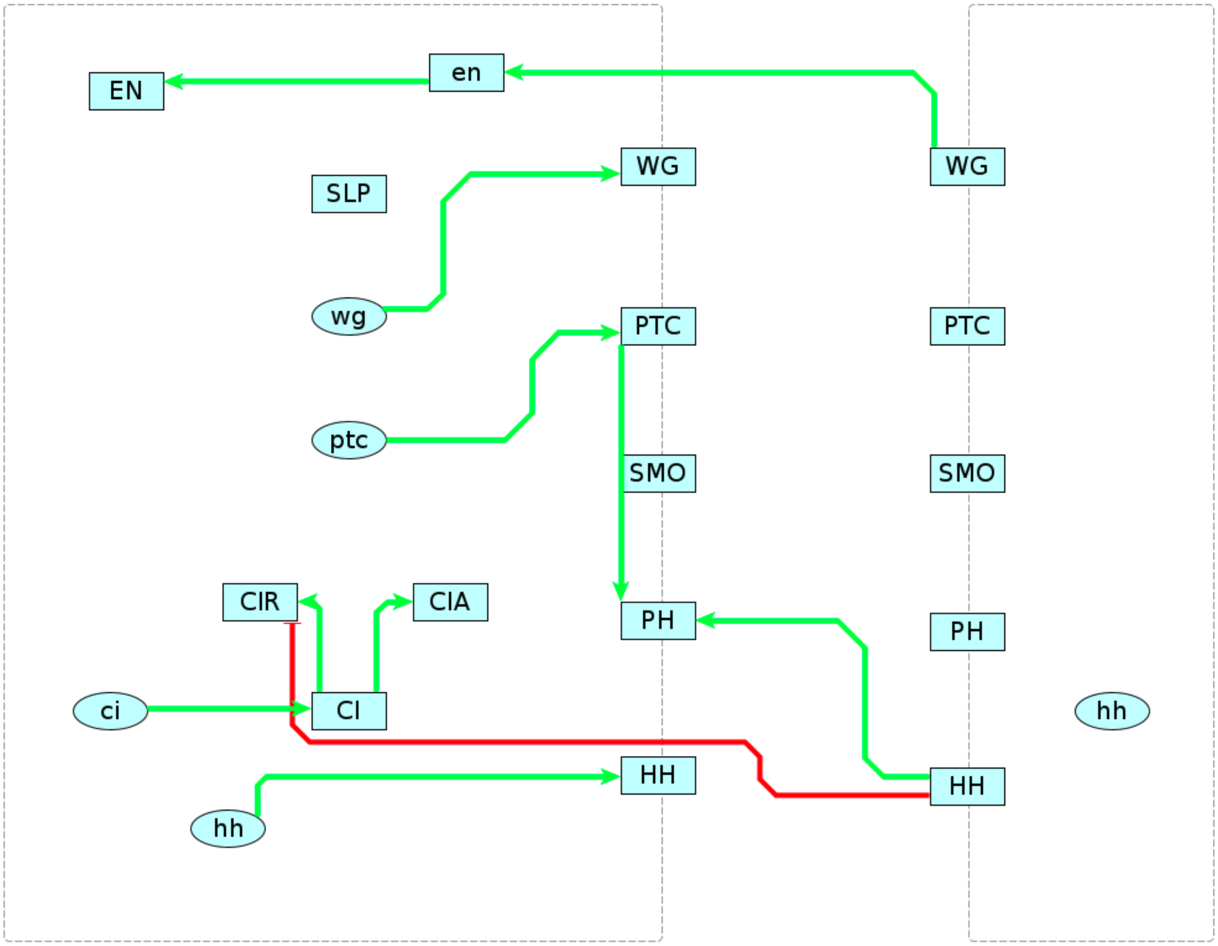
